# Design and Mathematical Analysis of Synthetic Inhibitory Circuits that Program Self-Organizing Multicellular Structures

**DOI:** 10.1101/2023.11.18.567649

**Authors:** Calvin Lam

## Abstract

Bottom-up approaches are becoming increasingly popular for studying multicellular self-organization and development. In contrast to the classic top-down approach, where parts of the organization/developmental process are broken to understand the process, the goal is to build the process to understand it. For example, synthetic circuits have been built to understand how cell-cell communication and differential adhesion can drive multicellular development. The majority of current bottom-up efforts focus on using activatory circuits to engineer and understand development, but efforts with inhibitory circuits have been minimal. Yet, inhibitory circuits are ubiquitous and vital to native developmental processes. Thus, inhibitory circuits are a crucial yet poorly studied facet of bottom-up multicellular development. To demonstrate the potential of inhibitory circuits for building and developing multicellular structures, I designed several synthetic inhibitory circuits that combine engineered cell-cell communication and differential adhesion. Using a previously validated *in silico* framework, I examine the capability of these circuits for synthetic development. I show that the designed inhibitory circuits can build a variety of patterned, self-organized structures and even morphological oscillations. These results support that inhibitory circuits can be powerful tools for building, studying, and understanding developmental processes.

**Graphical Abstract:** 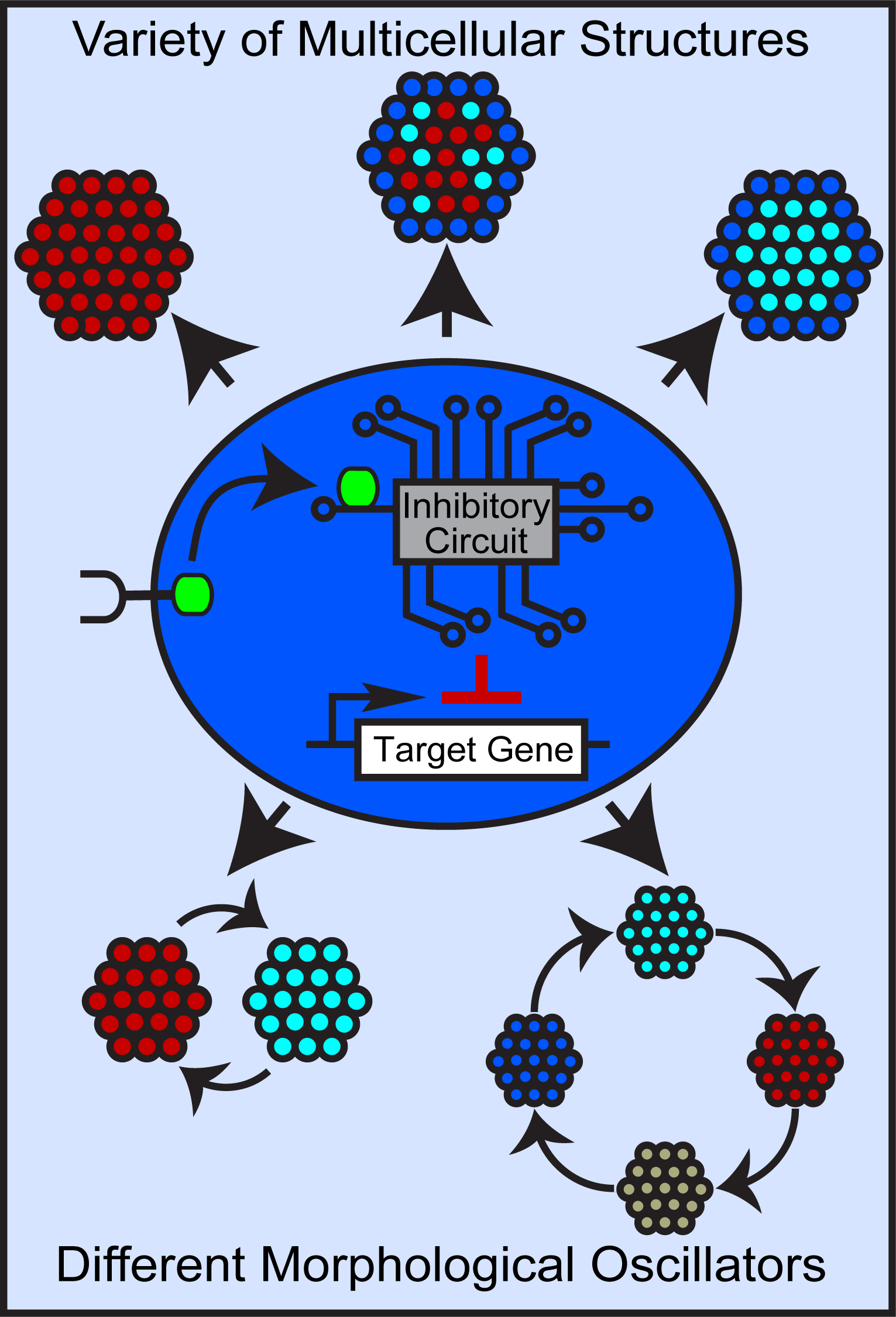

## INTRODUCTION

The development of multicellular organisms is an intricate, highly coordinated dance that has fascinated scientists for centuries ^1–9^. With minimal external control, individual units communicate with one another, alter their behavior accordingly, and self-organize into ornately patterned, functional structures ^10–17^. Understanding these developmental processes is a longstanding goal of biology; it not only provides insight into multicellular life, but also insight for clinical applications such as tissue engineering and regenerative medicine ^10,11,13–28^.

However, understanding these multicellular developmental processes is notoriously difficult. The classic top-down “break-it-to-understand-it” approach focuses on breaking a part of the process to understand the process, but breaking a part can affect the subsequent and parallel parts ^15,16,29^. This approach informs of the necessity of the part, but not necessarily the function(s) of the part ^16^.

In recent years, a complementary strategy has emerged through the field of synthetic biology. The field’s modular tools allow programming and controlling cells, enabling a bottom-up “build-it-to-understand-it” approach. In contrast to the top-down approach, the bottom-up approach seeks to join together parts that can construct/build the process, thereby providing understanding of the process ^10,11,13,14,16–23,25–27,29,30^. Though nascent, this approach has proven powerful thus far ^10,31^. Using synthetic juxtacrine receptors as parts to control cell-cell signaling processes, Morsut et al. demonstrated that such processes can build complex multicellular multi-layered patterns ^31^. Adding differential adhesion as a part illustrated that processes combining juxtacrine signaling with differential adhesion can build a variety of 3D multicellular patterned structures ^10^. These constructed processes provide insight into how basic components such as cell signaling and adhesion expression can direct complex self-organization. Moreover, these synthetic processes have features such as regeneration, cell fate divergence, and symmetry breaking, providing insight into how these features occur in native multicellular developmental processes ^10,31^.

The majority of these bottom-up, synthetic development efforts emphasize activation. For instance, the above works used a synthetic juxtacrine receptor to activate expression of a target gene (Fig.1A) ^10,31^. With different fluorescent reporter and adhesion cadherin genes as the target gene, these activatory circuits, when programmed into cells, yield developmental processes that can build a variety of patterned, multicellular self-organizing structures (Fig.1A) ^10,31^.

**Figure 1.**
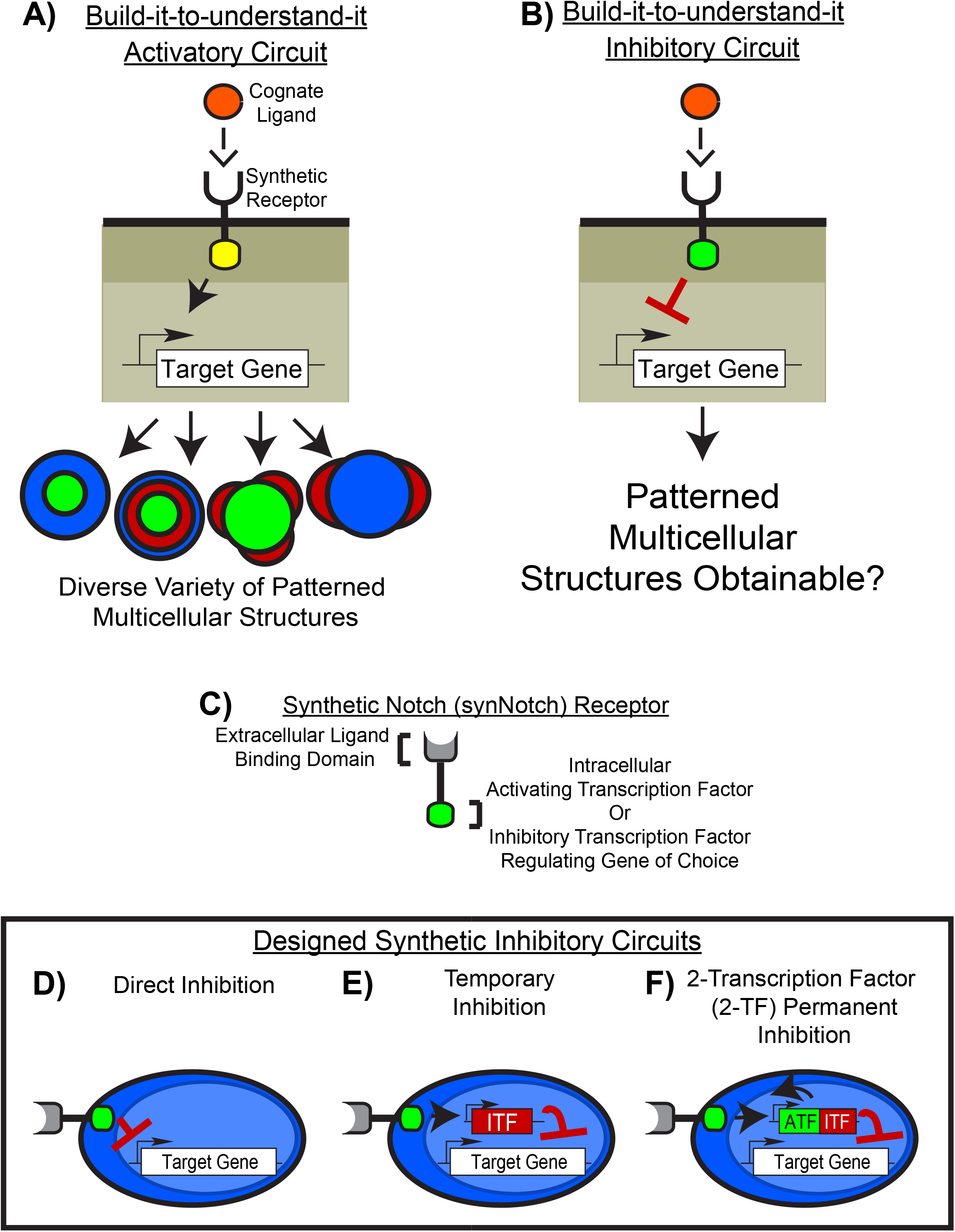
Design of the synthetic inhibitory circuits for bottom-up multicellular development. A) The majority of bottom-up “build-it-to-understand-it” approaches for multicellular development use activatory circuits. A target gene is activated to drive development. Such circuits are capable of building multitudes of structures. B) Inhibitory circuits, in contrast, are rarely used in the bottom-up approach. Are they also capable of building various multicellular structures? C) The synthetic Notch (synNotch) receptor is a synthetic receptor modular in both ligand binding and intracellular domain. Ligand binding domain allows sensing a ligand of choice and intracellular domain allows releasing an activating or inhibitory transcription factor of choice. D) The direct inhibition circuit has the synNotch receptor release an inhibitory transcription factor to directly repress target gene expression ^31^. E) The temporary inhibition circuit has the synNotch receptor first drive an inhibitory transcription factor (ITF) that then represses target gene expression. This circuit should result in temporarily prolonging gene repression. F) The 2-TF permanent inhibition circuit has the synNotch receptor drive a transgene cassette with an activating transcription factor (ATF) and ITF. ATF drives itself and the ITF, permanently expressing ITF to permanently repress gene expression.

To date, inhibitory circuits, despite being integral to native multicellular developmental processes, have been much less used in synthetic development (Fig.1B). Nonetheless, the few current results support that inhibitory circuits could be a powerful strategy for building and understanding multicellular development. For instance, using inhibitory circuits to build morphogen gradients revealed that double negative signaling logic coupled with negative feedback improves gradient pattern formation in the Sonic Hedgehog pathway ^14^. Using inhibitory circuits to inhibit morphogen activity allows the formation of sharp gradient boundaries ^11^. Inhibitory circuits coupled with differential adhesion can drive the formation of at least one type of patterned 3D structure ^10^.

To investigate the potential of inhibitory circuits for bottom-up multicellular development, I designed several generalizable synthetic inhibitory circuits. These circuits are driven by the synthetic Notch (synNotch) receptor, a powerful, fully modular receptor that is capable of both activating and inhibiting gene expression ^31,32^. Upon binding the juxtacrine cognate ligand, this receptor releases a transcription factor controlling the gene of choice (Fig.1C) ^31,32^. For gene activation, an activating transcription factor is released to drive target gene expression. For gene inhibition, the receptor can either directly repress gene expression by releasing an inhibitory transcription factor ^31^ or indirectly by driving expression of an inhibitory transcription factor that then represses target gene expression ^10^. The flexibility of the synNotch receptor allowed me to design inhibitory circuits with varying temporal dynamics for building and developing multicellular structures (Fig.1D-F) ^18^. In the published accompanying study, I showed how activatory circuits with varying spatiotemporal control can be used for bottom-up multicellular development ^18^.

Moreover, use of the synNotch receptor allowed me to employ a previously validated computational strategy. This enabled me to thoroughly explore the developmental capabilities of these inhibitory circuits ^18,19^. In a previous work, my colleagues and I developed a mathematical and *in silico* approach for modelling synNotch circuits that drive self-organization ^19^. The Generalized Juxtacrine Signaling Model (GJSM) set of equations allows simple and intuitive modeling of synNotch circuits. SynNotch circuits are first converted to GJSM equations and then implemented into *in silico* cells (via a framework such as the cellular Potts model ^33–35^). This computational approach can successfully predict the developmental structures that result from programming *in vitro* cells with synNotch circuits ^19^. This computational approach is ideal for exploring these circuits’ capabilities for bottom-up multicellular development; I can rapidly and systematically investigate circuit behavior across different parameters ^18^.

Here, I show that the designed inhibitory circuits are capable of their intended behavior. More importantly, when these circuits are combined with differential adhesion and implemented into *in silico* cells, these inhibitory circuits give rise to a variety of patterned multicellular structures. Of the structures obtained, the model predicts the only known *in vitro* patterned structure (expected as this was demonstrated in the original study ^19^), but the model also predicts that this only known structure is but a fraction of the morphologies possible. Further examination of the various structures indicated that one circuit is capable of morphological oscillations, but these oscillations dampen quickly, suggesting that further temporal regulation is required. Incorporating activating transcriptional amplifiers to additionally modulate temporal control revealed that the amplifiers can not only improve, but even rescue oscillations. These results support that inhibitory circuits can be powerful tools for bottom-up synthetic development.

## RESULTS

### Design of the synthetic inhibitory circuits

The direct inhibition circuit is the simplest of the designed inhibitory circuits, using a synNotch receptor that, upon binding its juxtacrine cognate ligand, releases a transcriptional repressor inhibiting expression of the target gene (Fig.1D) ^31^. Because repression is directly mediated by the synNotch receptor, this circuit is highly dependent on the presence of cognate ligand to maintain gene repression; loss of the signaling ligand should quickly result in decrease of gene repression ^18,31,32^. Thus, this design offers basic spatial control with minimal temporal control ^31,32^.

To demonstrate how inhibitory circuits can be used for bottom-up multicellular development, it would be ideal to also test other inhibitory circuits with additional levels of temporal control. I therefore designed two additional circuits: the temporary inhibition circuit and 2-transcription factor (2-TF) permanent inhibition circuit.

In the temporary inhibition circuit, I take advantage of the synNotch receptor’s modularity to also activate gene expression ^10,31,32^. The synNotch receptor drives expression of an inhibitory transcription factor (ITF) that then inhibits target gene expression (Fig.1E) ^10^. Repression is now dependent on the ITF level, rather than the synNotch signal, and thus gene repression should continue even when synNotch signaling is lost, as long as the ITF remains elevated enough to continue repression. Compared to the direct inhibition circuit, this circuit should enable temporarily prolonging gene repression.

In the 2-transcription factor (2-TF) permanent inhibition circuit, the synNotch receptor activates expression of both an activating transcription factor (ATF) and an inhibitory transcription factor (ITF) (Fig.1F). An example gene cassette would be the ATF and ITF genes linked by an internal ribosomal entry site or ribosomal skipping site, and expression controlled by a promoter activated by the synNotch receptor. The ATF drives itself and the ITF, providing a positive feedback loop for permanent ITF expression and thus should result in permanent gene repression ^18,36–38^.

### Circuits inhibit gene expression with temporal control over repression as designed

With the circuits designed, I converted them into GJSM equations and implemented them into *in silico* cells using the CompuCell3D cellular Potts framework ^33,34^. This combination was previously tested for modeling synNotch circuits for bottom-up multicellular development ^18,19^. See prior works ^18,19^ and the Methods section for more details. I then tested these circuits for their temporal control over gene repression. Blue cells, blue as they constitutively express blue reporter, were programmed to express a synNotch receptor that responds to orange ligand on orange cells (Fig.2A). In the direct inhibition circuit, the synNotch receptor directly inhibits blue reporter expression (Fig.2A). In the temporary inhibition circuit, the synNotch receptor drives an ITF that then inhibits blue reporter expression (Fig.2A). In the 2-TF permanent inhibition circuit, the synNotch receptor drives both an ATF and ITF. The ATF acts as a permanent amplifier of ITF expression and ITF inhibits blue reporter expression (Fig.2A) ^18^. I then tested these cells for their temporal control over gene repression using a simple cell-cell signaling setup from the accompanying study that examines activatory circuits ^18^.

**Figure 2.**
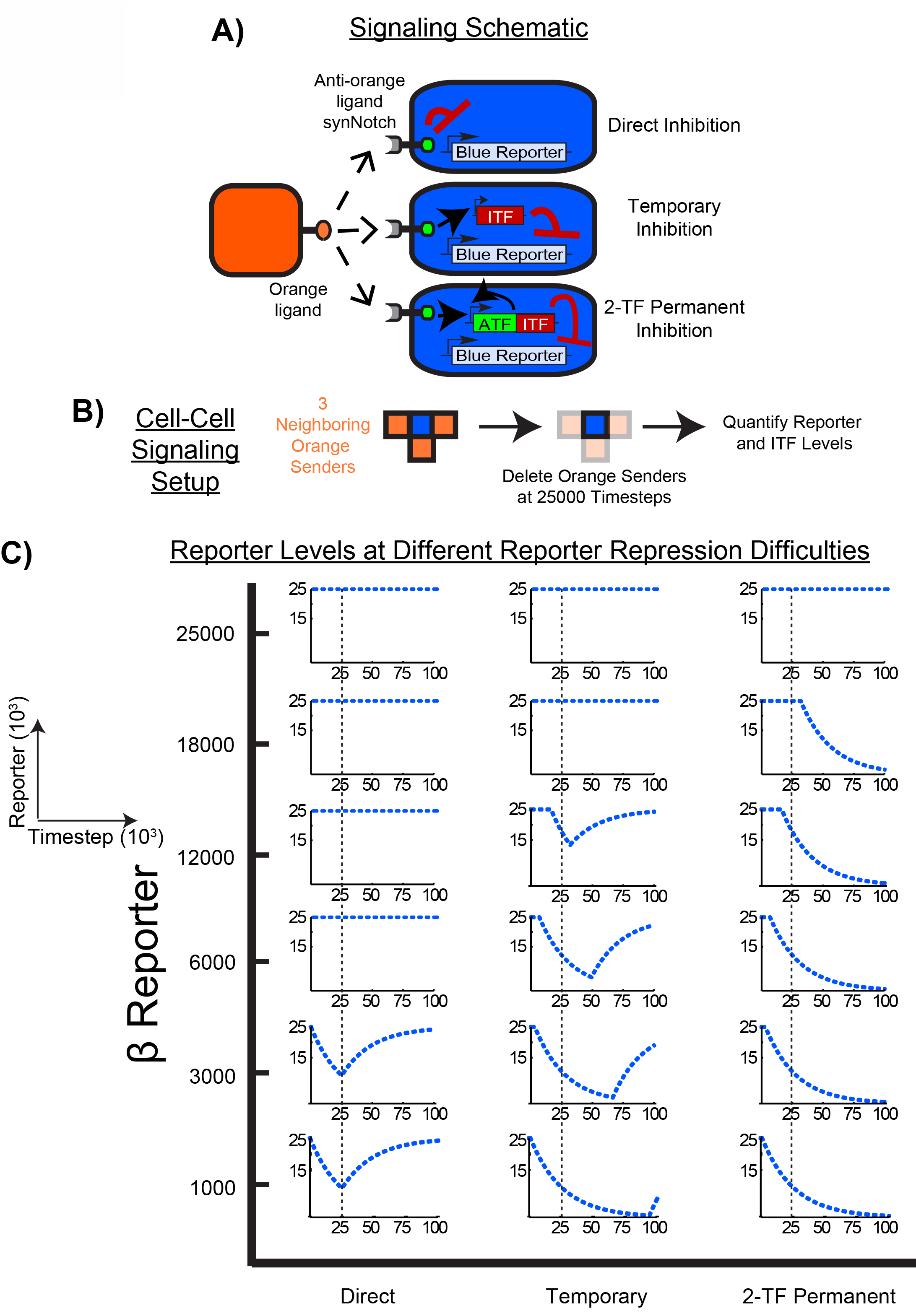
Inhibitory circuits inhibit gene expression as designed. A) Signaling schematic for testing if the inhibitory circuits operate as designed. Blue cells are programmed with an anti-orange ligand synNotch that triggers one of the inhibitory circuits to repress blue reporter expression. B) Cell-cell signaling test setup, where a blue cell is seeded with 3 orange neighbor cells and orange cells are deleted at 25000 timesteps to determine how orange ligand-synNotch signaling loss affects blue reporter repression. C) Blue reporter traces from the signaling test setup at different blue reporter repression difficulties (β reporter). Cells with the direct inhibition circuit lost gene repression immediately after orange neighbor loss, consistent with the circuit’s minimal temporal control design. These results are consistent with *in vitro* direct synNotch signaling dynamics ^18,31,32^. In contrast, the temporary inhibition circuit repressed at even higher repression difficulties and maintained repression for some time despite synNotch signaling loss. The 2-TF permanent inhibition circuit repressed at even further higher repression difficulties and maintained permanent blue reporter repression even after signaling loss. These results confirm that the circuits operate as designed. One trace shown per condition. Simulations run for 100000 timesteps.

A blue cell, imbued with one of the inhibitory circuits, is seeded with 3 orange sender cells. Cells are cubic and frozen (i.e. locked in shape, volume, and surface area) to maintain consistent synNotch signaling (Fig.2B). At 25000 timesteps, orange senders are deleted to test how ligand and synNotch signaling loss affects repression (Fig.2B). This setup allows controlling the exact level of synNotch signaling at a given time, thus enabling simultaneous testing of the circuit’s ability to inhibit reporter expression and temporality of repression ^18^.

In GJSM equations for synNotch circuits, gene expression and repression are modelled by several parameters. For gene activation or gene expression, β models gene expression difficulty while for inhibition or gene repression, β models gene repression difficulty. Higher values of β model higher expression/repression difficulties while lower values model lower expression/repression difficulties. Lower β values, however, do increase the risk of background gene expression/repression ^18,19^. κ models protein product degradation rate/saturation levels. See the accompanying study ^18^, the original study ^19^ or the Methods section for further details.

I therefore first checked the circuits for background gene inhibition. At reporter repression difficulties (β reporter) ≥1000, there was no background repression in all designed circuits (SFig.1A). I then tested the circuits at higher repression difficulties and found that all circuits were able to yield reporter repression (Fig.2C). The direct inhibition circuit was able to inhibit reporter expression up to β reporter=3000 and as predicted, had high spatial but minimal temporal control. Loss of synNotch signaling via loss of orange neighbors at 25000 timesteps resulted in immediate loss of blue reporter repression (Fig.2C). This temporal dynamic is consistent with reported synNotch signaling dynamics *in vitro* _18,31,32_.

In contrast, the temporary inhibition circuit repressed reporter expression up to β reporter=12000 and maintained inhibition for some time even after loss of orange neighbors (Fig.2C). Reporter levels eventually increased once again (Fig.2C) due to the ITF’s reliance on synNotch signaling to remain elevated (SFig.1B), confirming the design as a temporary inhibition circuit. The 2-TF permanent inhibition circuit repressed reporter expression up to β reporter=18000 and maintained permanent blue reporter repression even after loss of the orange neighbors (Fig.2C). This is due to the ATF’s positive feedback loop allowing permanent ITF expression and thus permanent reporter repression, confirming the circuit’s design as a permanent inhibition circuit (SFig.1B).

Similar results were obtained with 1 and 6 orange sender neighbors (data not shown). While all the inhibition circuits operate as designed, it is important to note that they fail at some parameter sets. Thus, circuit behavior is not solely defined by circuit design, but by its parameters as well ^18^.

### Temporary inhibition circuit can build a variety of patterned multicellular structures

With data supporting that the inhibitory circuits function as designed, I then tested the circuits’ for their ability to build and develop multicellular structures. A version of the temporary inhibition circuit, the lateral inhibition circuit, has been shown capable of driving bottom-up synthetic development *in vitro* ^10^. In this circuit, cells signal to one another via a CD19 synNotch interaction to drive expression of the transcriptional repressor tTs and the adhesion protein E-cadherin (gene is *ECAD*) (Fig.3A). tTs repressor then inhibits expression of the CD19 ligand. L929 mouse fibroblasts programmed with this circuit reliably form only one type of patterned structure, a 2-layered structure with CD19^-^ E-cadherin^+^ green cells and CD19^+^ E-cadherin^+^ yellow cells in the center (Fig.3B). CD19^+^ E-cadherin^-^ red cells form the peripheral ring (Fig.3B).

**Figure 3.**
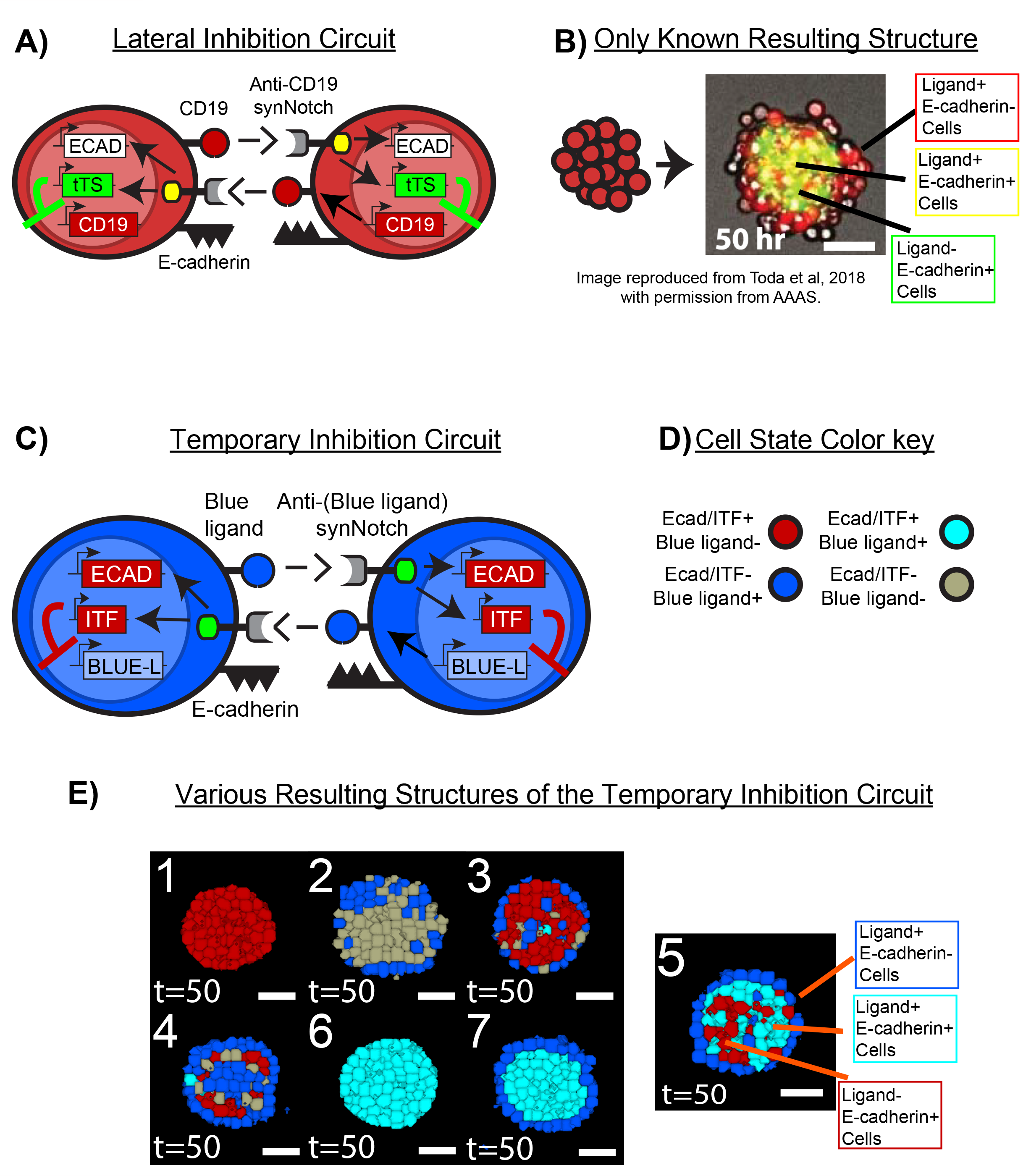
Temporary inhibition circuit can build a variety of multicellular structures. A) One version of the temporary inhibition circuit, the lateral inhibition circuit ^10^. In this version, anti-CD19 synNotch drives E-cadherin (*ECAD*) and tet transcriptional repressor *(tTS*) that represses CD19 expression. B) Mixing ∼100 cells programmed with this circuit results in a 2-layered structure with CD19^-^ E-cadherin^+^ green cells and CD19^+^ E-cadherin^+^ yellow cells in the center with CD19^+^ E-cadherin^-^ red cells on the periphery. Micrograph reproduced from Toda et al, 2018 ^10^ with permission from AAAS. C) The temporary inhibition circuit is the generalized version of the lateral inhibition circuit. ECAD is E-cadherin. ITF is generic representation for tTs. BLUE-L (blue ligand) is generic representation for CD19. D) A cell state color key is given to match expression state to cell color in the subsequent images. E) Mixing 93 blue cells programmed with the temporary inhibition circuit results in multitudes of patterned structures. A representative set of structures is given, showcasing the variety of patterned structures that can form from the temporary inhibition circuit. Each structure resulted from a different parameter set of expression difficulties for ITF/ECAD and repression difficulties for blue ligand. The full gallery of structures, along with the parameters that generated them, is given in SFig.2A. As expected, and previously shown, the model predicts the known *in vitro* structure ^19^. These results indicate that the only known structure to date is a small subset of structures possible with this circuit. N=3 for each structure. Simulations run for 50000 timesteps.

The robust formation of the 2-layered structure is likely due to the lateral inhibition circuit being partially calibrated ^10^, thereby limiting the structures buildable. However, systematically investigating the generalized version of the lateral inhibition circuit, the temporary inhibition circuit, revealed that this circuit design can form a wealth of patterned structures (Fig.3E and SFig.2A). The circuit was implemented in *in silico* L929 (ISL929) cells ^18,19^ to parallel the *in vitro* implementation. I tested a wide range of expression difficulties for ITF/ECAD (ITF is generic representation of tTS, gene representation is *ITF*) and repression difficulties for blue ligand (blue ligand is generic representation of CD19, gene is *BLUE-L)*. A cell color guide is given in Fig.3D. Scanned parameters are in SFig.2A.

Several structure types were formed, ranging from types 1 and 6 of highly homogenous E-cadherin^+^ spheroids to 3, 4, 5, and 7 for 2-layered structures to 2 for mixed E-cadherin^-^ spheroids (Fig.3E). Representative structure types are shown and additional structures, organized by the parameters that generated them, are given in SFig.2A. Out of all the structure types obtained, only one (type 5 of Fig.3E), was the 2-layered structure built by the lateral inhibition circuit (Fig.3B). These results indicate that the temporary inhibition circuit can build a variety of patterned, multicellular structures. The only known *in vitro* example is but a fraction of the structures possible.

### Direct inhibition circuit and 2-TF permanent inhibition circuit can also build a variety of patterned multicellular structures

Testing the direct inhibition circuit and 2-TF permanent inhibition circuit revealed that both circuits can also generate a variety of patterned structures. The direct inhibition circuit built structure types ranging from types 1 and 3 of heterogenous E-cadherin^+^ spheroids to 4, 6, and 7 for 2-layered structures to 2 and 5 for E-cadherin^-^ spheroids (Fig.4A). The full gallery of structures, organized by the parameters that generated them, is given in SFig.2B.

**Figure 4.**
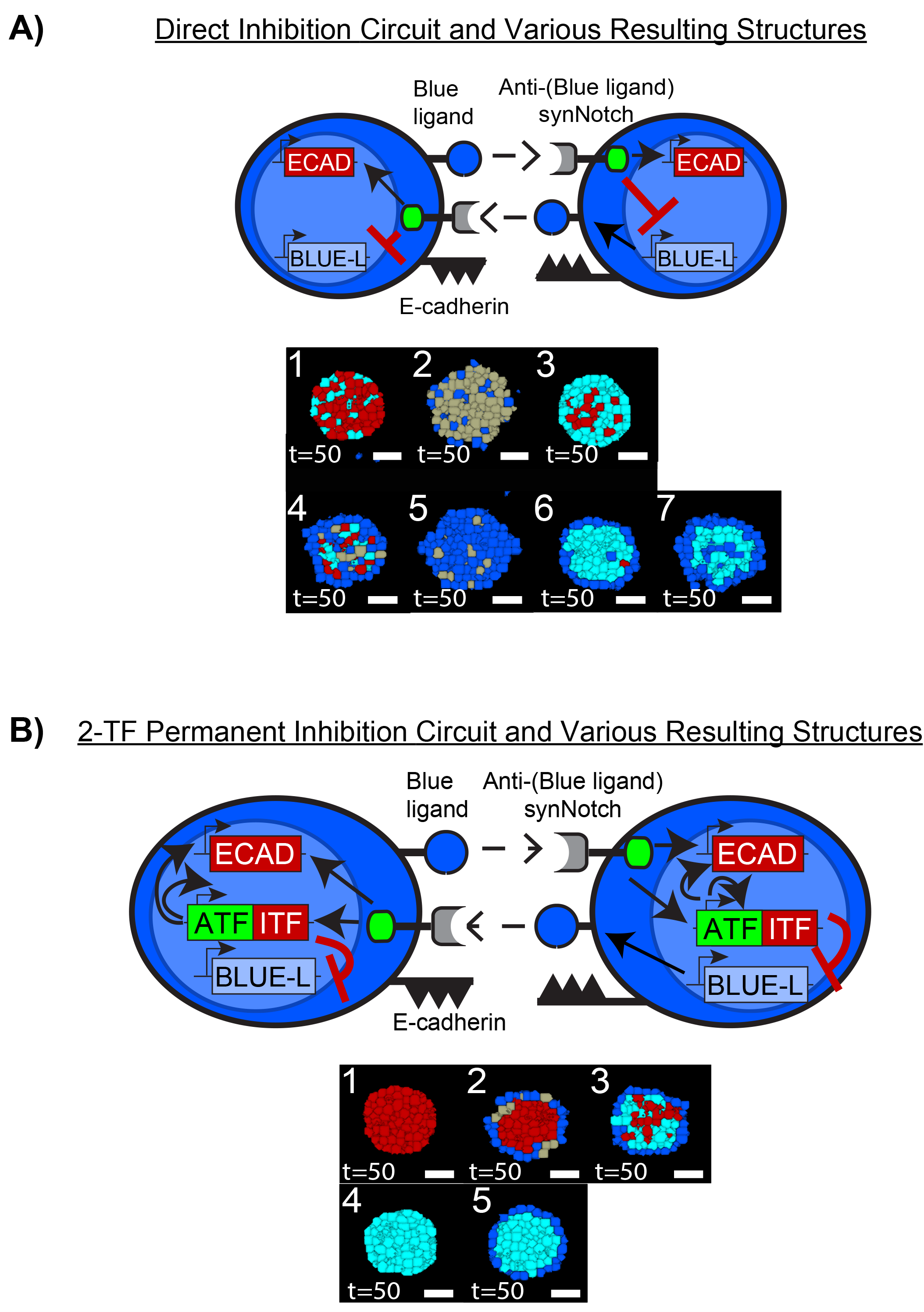
Designed inhibitory circuits can build multitudes of multicellular structures. A) The direct inhibition circuit can also build various patterned structures. 93 blue cells with this circuit results in 7 different types of patterned structures. One representative structure for each pattern type is shown. Full gallery of structures is given in SFig.2B. B) The 2-TF permanent inhibition circuit can also build various patterned structures. 93 blue cells with this circuit results in 5 different types of patterned structures. One representative structure for each pattern type is shown. Full gallery of structures is given in SFig.2C. N=3 for each structure. Simulations run for 50000 timesteps.

Likewise, the 2-TF permanent inhibition circuit also proved capable of building various patterned structures. Structure types ranged from 1 and 4 for highly homogenous E-cadherin^+^ spheroids to 2, 3, and 5 for 2-layered structures (Fig.4B). Interestingly, this circuit yields less structure types compared to the other two, possibly due to its permanency in gene repression. The full gallery of structures, organized by the parameters that generated them, is given in SFig.2C.

Altogether, these results (Figs.3-4, SFig.2) demonstrate that inhibitory circuits can be powerful tools for building multicellular structures.

### Temporary inhibition circuit can build oscillatory structures

Of the diverse structures built by the inhibitory circuits, the homogenous spheroids of the temporary inhibition circuit (structures of types 1 and 6 of Fig.3E) were of notable interest. During the development of these structures, cells were highly uniform in the expression state of ITF/E-cadherin and blue ligand (i.e. all cells were ITF/E-cadherin^+^ blue ligand^+^ or ITF/E-cadherin^+^ blue ligand^-^ etc, see Fig.3D for all possible expression states and cell color for each state). Cells then synchronously transitioned from one expression state to another expression state (i.e. cells transitioned from one color to another color at a similar time). An example development with these features is shown in Fig.5A. These features are strikingly similar to those observed with oscillations in development ^6,35,39,40^. Because temporary gene repression is known to drive oscillation ^6,35,39–43^, I hypothesized that the temporary inhibition circuit can build morphological oscillators (Fig.5B).

**Figure 5.**
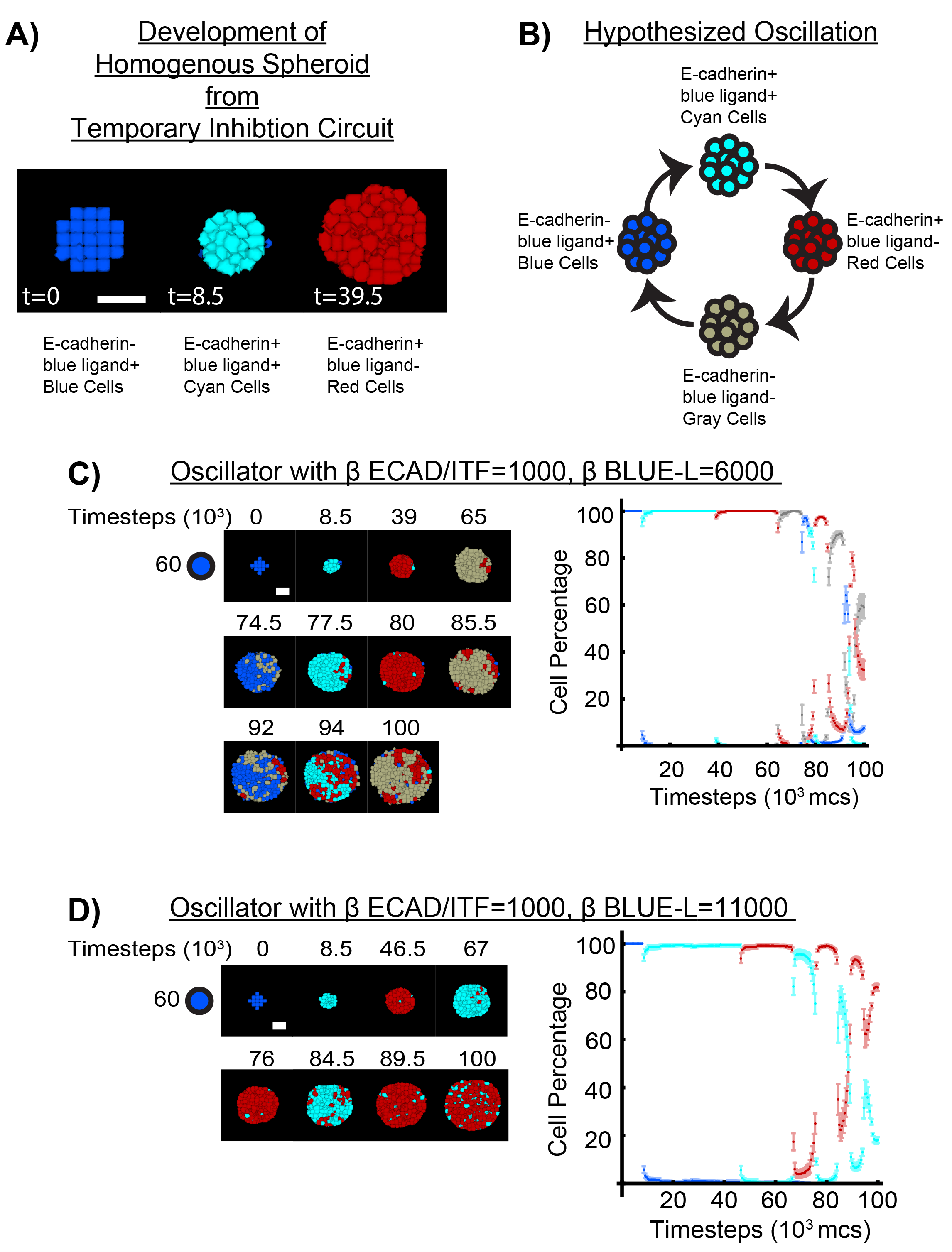
Temporary inhibition circuit can build oscillatory structures. A) Homogenous spheroids built by the temporary inhibition circuit had cells synchronized in expression state (color) with sharp transition to the next expression state. An example developmental trajectory with these synchronous and sharp transitions is shown. Cell expression state is given below each image. B) This led me to hypothesize that the temporary inhibition circuit can build oscillatory structures. Hypothesized oscillation is shown along with cell expression state for each phase in the oscillation. C) 57 blue cells programmed with the temporary inhibition circuit and ECAD/ITF expression difficulty=1000 (β ECAD/ITF=1000) with blue ligand repression difficulty=6000 (β BLUE-L=6000) generated morphological oscillation. Cells synchronously transitioned from blue to cyan to red to gray before repeating the oscillation. Developmental trajectory shown is of images at the transition time. Plot of cell percentage is given as well. D) 57 blue cells programmed with the temporary inhibition circuit but β ECAD/ITF=1000 and β BLUE-L=11000. Oscillation differed from that of C, skipping the blue and gray phase. Developmental trajectory is shown along with plot of cell percentage. Additional oscillations with different parameters are given in SFig.3. A representative developmental trajectory is given per oscillation. N=3 for each oscillation. Simulations run for 100000 timesteps.

Extending the simulation time to 100000 timesteps for the pink outlined structures of SFig.2A revealed that the temporary inhibition circuit can indeed build morphological oscillators (Fig.5C-D, SFig.3). An initial structure of blue cells sharply transitions to cyan before transitioning to red and then gray before repeating the cycle (Fig.5C). Testing the circuit with a higher blue ligand repression difficulty β BLUE-L=11000 built a morphological oscillator that skipped the gray and blue phases, yielding a cyan red oscillator (Fig.5D). Oscillators with the remaining parameters (β BLUE-L=16000, 21000) are given in SFig.3. All these oscillations were highly reproducible and consistent (n=3 each). β BLUE-L=1000 was a notably poor oscillator (SFig.3C). With the same parameters, the direct inhibition circuit and 2-TF permanent inhibition circuit did not yield oscillatory structures, confirming that temporary gene repression is the driver of oscillation (SFig.4) ^6,35,39–43^.

### Additional temporal control via activating amplifiers improves oscillation quality

In addition to being able to build multicellular structures, inhibitory circuits can also generate morphological oscillations, demonstrating their potential for bottom-up synthetic development. It would be even more powerful, however, if the oscillations could be further modified and improved. The morphological oscillations generated by the temporary inhibition circuit dampened quickly (Fig.5C-D). These results were confirmed by running the simulations for longer (SFig.5A,C).

In the accompanying study ^18^, I showed how activatory circuits, through activating transcriptional amplifiers, can be used to enact spatiotemporal control over gene expression in bottom-up multicellular development. As temporary gene repression is what drives oscillation, I reasoned that using an activating amplifier to control the temporary gene repression of the temporary inhibition circuit could improve oscillation (Fig.6A). The circuit design (Fig.6B) is a simple modification of the temporary inhibition circuit; it uses an activating amplifier to control the inhibitory circuit of Fig.3C. The synNotch receptor now drives an activating amplifier consisting of an activating transcription factor (ATF) (Fig.6B). The ATF then drives expression of E-cadherin and ITF (Fig.6B). ITF then inhibits blue ligand expression as in the original circuit (Fig.6B).

**Figure 6.**
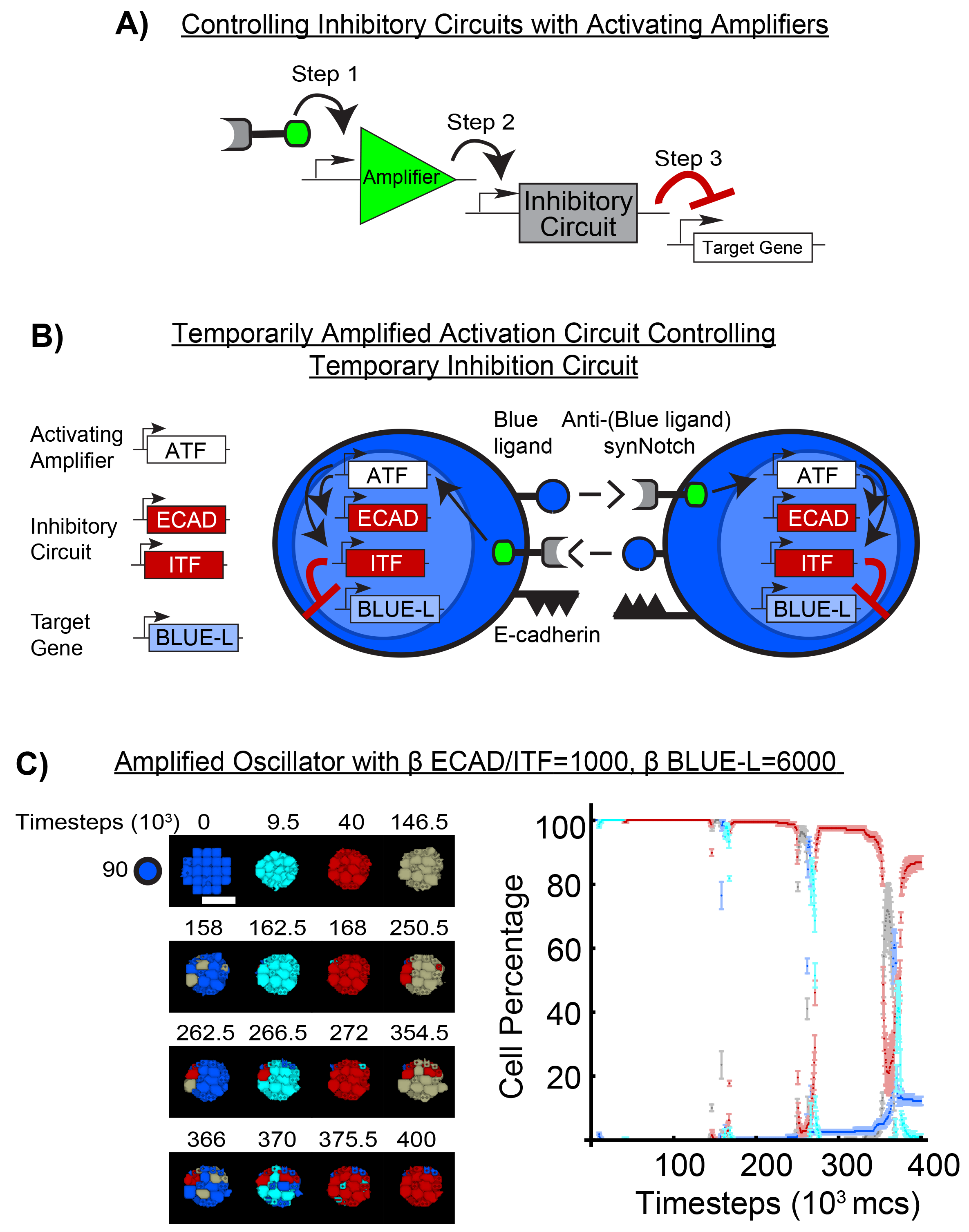
Using activating amplifiers to control the temporary inhibition circuit can improve oscillation quality. A) I recently showed that activating amplifiers can modulate temporal dynamics in synthetic circuits ^18^. I therefore tested if prolonging repression with an activating amplifier could improve oscillation quality. B) Temporarily amplified activation circuit controlling temporary inhibition circuit. The temporary inhibition circuit is rewired such that the synNotch receptor now drives the activating amplifier (activating transcription factor, ATF). The activating amplifier then drives the inhibitory circuit that then represses target gene expression. C) Mixing 93 blue cells programmed with this circuit builds an oscillator with improved oscillation quality compared to its non-amplified variant (SFig.5A). Mitosis was removed due to computational limitations, but this does not affect timing or quality of oscillation (compare Fig.5C to SFig.5A, Fig.5D to SFig.5C). Developmental trajectory is shown along with plot of cell percentage. Additional oscillations with different parameters are given in SFig.5-7. A representative developmental trajectory is given per oscillation. N=3 for each oscillation. Simulations run for 400000 timesteps.

Using the same parameters as the temporary inhibition circuit oscillator of Fig.5C and SFig.5A, the amplified oscillator improved morphological oscillation (Fig.6C). I compared oscillation quality between circuits using the number of cyan phases completed before half amplitude (ring-down method in signal analysis) ^44^. I chose the cyan phase for quantification as it is present in all oscillators (i.e. cyan red oscillators lack gray and blue phases (Fig.5D)) and does not have constant drift (i.e. see SFig.7B).

Compared to its non-amplified variant (Fig.5C and SFig.5A), the amplified oscillator (Fig.6C) was approximately a 1.5-fold improvement in oscillation quality (3 cyan phases to 2 cyan phases). Due to prolonged inhibition from the activating amplifier ^18^, oscillation took notably longer, requiring almost 400000 timesteps for the three cycles (Fig.6C). Dampening was also observed but is notably less prominent. The oscillator of SFig.5A had red cell percentage quickly drift up with each red phase (5% to 20% to 50%) (SFig.5A). In contrast, the amplified oscillator had a much lower drift (0% to 5% to 20%) (Fig.6C).

Similar results were obtained for the β ECAD/ITF=1000 β BLUE-L=11000 amplified oscillator (SFig.5B). The amplified oscillator was at least a 1.3-fold improvement in oscillation quality compared to its non-amplified counterpart (SFig.5C) (4 cyan phases to 3 cyan phases). The oscillation occurred over a longer duration due to prolonged inhibition from the activating amplifier as expected (SFig.5B). Dampening was observed but the amplified oscillator only had red percentage drift from 5% to 10% to 20% in the observed red phases (SFig.5B) while in the equivalent red phases of the oscillator, drift was from 0% to 40% to 70% (SFig.5C).

To confirm that oscillations are not limited to β ECAD/ITF=1000 circuits, I tested if β ECAD/ITF=6000 circuits could also generate oscillatory structures. Oscillators with β ECAD/ITF=6000 and β BLUE-L=1000 or 6000 generated no oscillations (SFig.6A and SFig.7A). However, the amplified oscillator circuits were able to oscillate (SFig.6B and SFig.7B). These results indicate that incorporating activating amplifiers can not only improve, but even rescue oscillations.

These results, in conjunction with those of Fig.5 and SFig.3, indicate that inhibitory circuits are powerful tools for achieving morphological oscillations. These circuits can be flexibly modified with additional circuits to alter their oscillatory behavior, supporting their utility for bottom-up synthetic development.

## DISCUSSION

The bottom-up approach is an emerging but powerful strategy for studying multicellular development. Most bottom-up efforts focus on activatory circuits to drive synthetic development, but in this study, I show that inhibitory circuits can also be powerful tools for driving development. Via an *in silico* approach, I show that the designed inhibitory circuits can build a variety of patterned multicellular structures. A systematic parameter scan reveals that the only known inhibitory structure to date is but a fraction of the structures possible with inhibitory circuits. Further examination of one circuit revealed that it is capable of morphological oscillations and that these oscillations could be improved by using activatory circuits from the accompanying study. Altogether, these results support that inhibitory circuits can be powerful tools for building and studying developmental processes.

To demonstrate how inhibitory circuits can be used to build multicellular structures, I employed a previously tested computational approach ^18,19^. By combining the GJSM with a cellular Potts framework, I was able to test the circuits across a variety of parameter combinations and show the numerous different morphologies buildable from these circuits (Fig.3-4, SFig.2) ^18,19,34^. This computational setup has been validated for modelling and predicting the self-organizing structures that arise from synNotch circuits driving differential adhesion ^19^. Nonetheless, I recognize that a key limitation of computational studies is experimentational validation. Unfortunately, as an independent investigator, I lack all the fundings and resources to perform any of the analogous biological experiments. With this limitation in mind, at the inception of this study, I deliberately designed and chose to test the direct inhibition circuit and temporary inhibition circuit as specific biological versions of these circuits exist ^10,31^. The specific version of the direct inhibition circuit uses Gal4-KRAB as the inhibitory transcription factor ^31^. Its inhibition and temporal dynamics are consistent with the results obtained for the direct inhibition circuit in Fig.2C. The specific version of the temporary inhibition circuit, the lateral inhibition circuit (Fig.3A), served as an additional check for my strategy. The lateral inhibition circuit’s only known structure was predicted by the *in silico* approach (Fig.3E), confirming the results of the previous study ^19^ and supporting the validity of the computational approach. The *in silico* approach additionally predicted a variety of other structures yet to be obtained *in vitro* (Fig.3E), but this reflects a difference in methodology. The *in vitro* lateral inhibition circuit was partially calibrated ^10^, thereby limiting the types of structures buildable, while the *in silico* temporary inhibition circuit was systematically examined across numerous different parameters.

In addition, the *in silico* model also predicts that the temporary inhibition circuit is capable of morphological oscillations (Fig.5, SFig.3). Temporary repression is well-known to drive oscillations in single cells, but it has yet to be shown how oscillations in multiple cells can drive synthetic multicellular development ^41–43,45,46^. This result would be particularly interesting to test biologically, as it could demonstrate that inhibitory circuits can build dynamic structures as well. Such results would indicate that inhibitory circuits are indispensable tools for bottom-up synthetic development.

As in the accompanying study ^18^, I designed the circuits generically to accommodate the rapidly increasing toolkit of synthetic biology ^31,38,47–62^. I leave the component choice to the user so that future components can be incorporated and tested. For example, typical transcription factors (TFs) at the time of the lateral inhibition circuit included the ATFs Gal4-VP64, LexA-VP64, tTa and ITF Gal4-KRAB ^31,32,38,63–65^. Since then, however, new TFs have emerged with improved compatibility and modularity ^47–49,66^.

These TF development efforts, like most bottom-up efforts, notably focus on gene activation. For bottom-up approaches to advance, there needs to be efforts towards developing new and improved ITFs as well. As an example, the direct inhibition circuit that builds multicellular structures (Fig.4A) requires a bifunctional transcription factor that can both inhibit and activate gene expression. Such a transcription factor has yet to be engineered and used for multicellular synthetic development, but natural bifunctional transcription factors do exist and could be a viable starting point for this component ^67–70^.

Though the circuits here are relatively basic in design, they can build a wealth of patterned multicellular structures and even morphological oscillations. They even have features of native developmental processes such as cell fate divergence, self-organization, and synchronized oscillation. “Build-it-to-understand-it” efforts with inhibitory circuits could advance our understanding of native multicellular developmental processes. Beyond multicellular development, inhibitory circuits, like activatory circuits, could potentially be used for therapeutic purposes as well such as regenerative medicine and synthetic organ development ^18,21,23,24^. This work supports future research on inhibitory circuits, and I hope it will promote the use of inhibitory circuits in multicellular synthetic biology.

## ACKNOWLEDGEMENTS

I would like to thank UNMC for access to and aid in printing research articles. I am thankful to the three anonymous reviewers of my previous article ^18^ as their suggestions have also helped improve this manuscript.

## AUTHOR CONTRIBUTIONS

C.L. conceived, designed, performed, analyzed, and wrote the entire study.

## METHODS

### Key Resource Table

| SOFTWARE AND ALGORITHMS | SOURCE | IDENTIFIER |
| --- | --- | --- |
| CompuCell3D (CC3D) v3.7.8 | <sup>34</sup> | RRID:SCR_003052 |
| Mathematica v12.0.0.0 | Wolfram Research | RRID:SCR_014448 |
| Excel v2310 | Microsoft | RRID:SCR_016137 |
| General Juxtacrine Signaling Model (GJSM) | <sup>19</sup> | N/A |
| Activating Transcriptional Amplifiers in GJSM | <sup>18</sup> | N/A |

### Lead Contact

Requests for information and code should be directed to Calvin Lam (<u>calvin.lam.k@gmail</u>).

### *In silico* cell line ISL929

ISL929 is an *in silico* cell line developed in ^19^ to model the *in vitro* L929 cell line used for building multicellular structures with the synNotch receptor ^10^. This line was developed in the CompuCell3D cellular Potts framework ^34^ and has been used with the Generalized Juxtacrine Signaling Model (GJSM) to successfully predict how synNotch circuits can drive multicellular development ^19^.

Because the lateral inhibition circuit was implemented in L929 cells ^10^, ISL929 served as the ideal *in silico* cells for this study. It would allow me to check the previous study’s results ^19^ while also providing a biological experiment start point such as testing the inhibitory circuits from this study in L929 cells. A brief description of ISL929 is given in the accompanying study ^18^ and a complete description is in the original study ^19^.

### Overview of the Generalized Juxtacrine Signaling Model

The Generalized Juxtracine Signaling Model (GJSM) was developed to model synthetic juxtacrine receptor (i.e. synNotch ^10,31^, SNIPR ^62^) regulation of gene expression ^19^. When parameterized, the model can capture synthetic juxtacrine receptor signaling dynamics and has successfully predicted the developmental structures that arise from synNotch-based differential adhesion circuits ^19^. A brief description of the relevant model features is given below. Full descriptions can be found in ^18,19^.

Gene expression by a synthetic juxtacrine receptor such as synNotch can be described by the general equation

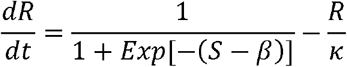

The change in the expressed gene’s protein level R at a given timestep t is a function of both production (first term) and degradation (second term). In the production term, signal S models receptor or activating transcription factor (ATF) signaling driving gene expression. β models gene expression difficulty and encompasses the biological processes that affect gene to functional protein product such as such as promoter, transcription, translation, trafficking inefficiencies, and expression delay, etc ^18,19^.

Degradation (second term) is modelled by the standard linear decay model. Protein product level R decays proportional to itself and inverse to decay constant κ. As a result, κ controls not only decay rate but saturation level as well.

The logistic form was chosen for GJSM over the Hill form for several reasons as described in the original study ^19^ and accompanying study ^18^. First, the logistic form has simple and intuitive parameter interpretations that allow a user new to modeling to easily start. Weighing S against β allows a user to quickly understand how expression difficulty and signal affects expression. If S largely exceeds expression difficulty, then expression is easy. If expression difficulty largely exceeds S, then expression is difficult. Second, because the logistic form is mathematically equivalent to the Hill form, a more advanced user can convert the logistic equations to Hill equations to obtain more biologically relevant parameters if desired ^18,71,72^.

Because the logistic function is equivalent to the Hill function, GJSM is also capable of modeling gene repression. This was shown in the original study ^19^. Gene repression in GJSM can be described by the equation

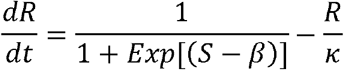

This form is mathematically obtained by transforming the expression equation with S to -S and β to – β ^19^. Signal S now models receptor or inhibitory transcription factor (ITF) signaling inhibiting gene expression. β now models gene repression difficulty and still encompasses the processes of gene to functional protein product as in the expression equation. Intuitive interpretations remain and degradation modelling remains the same as in the expression equation.

### Inhibitory circuits as modelled by GJSM

With the general equations described, equation sets for the inhibitory circuits can now be defined.

#### Direct Inhibition Circuit

In the direct inhibition circuit, the synNotch receptor directly inhibits target gene expression (Fig.1D). Then, the circuit equation is

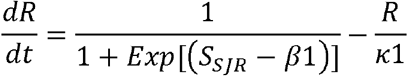

S_SJR_ is the number of activated synNotch receptors at a given timestep, with activated meaning bound to cognate ligand and released its transcription factor. Parameters are as described above for inhibitory circuits. How S_SJR_ is calculated is in a below section.

In the direct inhibition circuit for building multicellular structures (Fig.4A), the synNotch receptor also drives expression of the adhesion protein E-cadherin. E-cadherin levels are described by

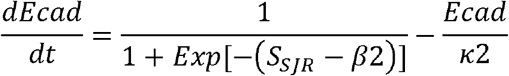

S_SJR_ is the number of activated synNotch receptors at a given timestep. Parameters are as described above for activatory circuits. How S_SJR_ is calculated is in a below section.

#### Temporary Inhibition Circuit/Oscillator Circuit

In the temporary inhibition circuit/oscillator circuit, the synNotch receptor indirectly inhibits gene expression by driving expression of an inhibitory transcription factor (ITF) that then inhibits target gene expression (Fig.1E). ITF levels are described by

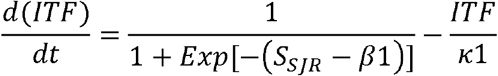

Parameters are as described above for activatory circuits. Target gene product level R is then described by the inhibitory equation

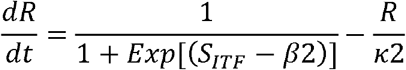

S_ITF_ is the ITF level at a given timestep from solving the first equation in the circuit. Parameters are as described above for inhibitory circuits.

In the temporary inhibition circuit for building multicellular structures (Fig.3) or oscillators (Fig.5), the synNotch receptor, in addition to driving the ITF, also drives expression of E-cadherin. To simplify the number of parameters and equations used per circuit, E-cadherin levels are described by the ITF equation.

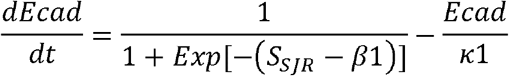

However, if desired, E-cadherin can be defined separately from ITF with its own parameters governing expression difficulty and decay.

#### 2-Transcription Factor (2-TF) Permanent Inhibition Circuit

In the 2-TF permanent inhibition circuit, the synNotch receptor drives expression of the activating transcription factor (ATF) and ITF (Fig.1F). The ATF is also able to drive expression of the ATF and ITF (Fig.1F). Then, ATF levels are described by

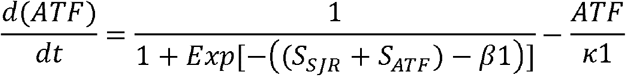

S_ATF_ is the ATF level from solving this equation at a given timestep. Parameters are as described above for activatory circuits.

To simplify parameters and equations used and because the ITF is linked to ATF expression, the ITF is assigned the same equation

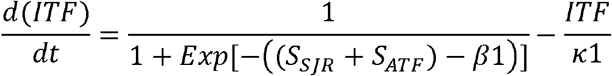

The ITF then inhibits target gene expression as in the temporary inhibition circuit. Thus, the equation is

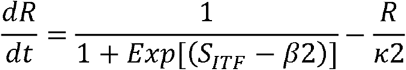

In the 2-TF permanent inhibition circuit for building multicellular structures (Fig.4B), in order to avoid the use of hybrid combinatorial promoters, which are not tested with GJSM ^18^, E-cadherin must be controlled by the same type of promoter as the ATF/ITF. Because ATF is able to drive expression of itself through this promoter, E-cadherin must also be driven by the ATF. E-cadherin is also driven by the synNotch receptor. Then, to simplify the number of parameters and equations used per circuit, E-cadherin levels are described by the ATF/ITF equation

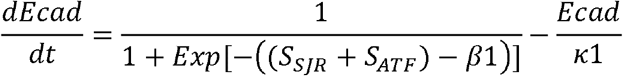

As in the temporary inhibition circuit, E-cadherin can be defined separately from ATF/ITF with its own parameters if desired.

#### Amplified Oscillator Circuit (Temporarily Amplified Activation Circuit Controlling Temporary Inhibition Circuit)

In the amplified oscillator circuit, a temporarily amplified activation circuit controls the temporary inhibition circuit (Fig.6B). The synNotch receptor drives expression of an ATF that then controls the temporary inhibition circuit.

The ATF equation is as in the published accompanying study ^18^

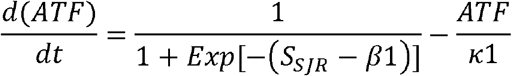

This ATF then drives expression of ITF and E-cadherin, replacing the role of the synNotch receptor in the temporary inhibition circuit. Then, the equations are

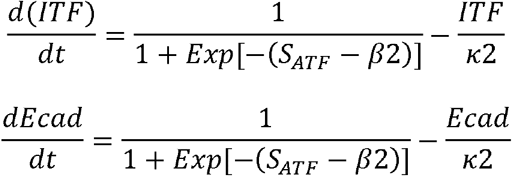

Finally, the ITF then inhibits target gene expression as in the temporary inhibition circuit. Thus, the equation is

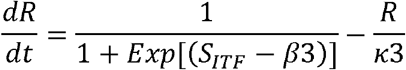

For the equations of each circuit, parameters that do not change between equations are denoted with the same variable (i.e. β1 is the same across all equations in the same circuit). However, it is important to note that parameters can be set differently between equations if desired. This confers additional flexibility in modeling, but increases the parameters used for a circuit. Parameters are given in Supplementary Table 1.

### Programming ISL929 cells with circuits and receptors

Having defined the equations for each circuit, I then implemented them into ISL929 cells. See SFig.1A of the accompanying study for a graphical depiction of this process ^18^. I then added the appropriate constitutive ligands into the appropriate cells (i.e. orange ligand on orange cells). I also added synNotch to the appropriate cells as well. To simplify calculations, I assumed synNotch to be in excess and non-limiting as in the reference and accompanying studies ^10,18,19^. If desired, the complete GJSM framework can be used to model receptor limiting cases. See the original study ^19^ for the receptor limiting formulation of GJSM. With these rules defined, it is now possible to calculate the signal S_SJR_ from synNotch signaling in the circuit equations. The relevant calculations are given here. The complete description and generalized formula can be found in the original study ^19^.

SynNotch functions in a 1:1 stoichiometry; one activated receptor releases one transcription factor regulating the target gene ^18,19,31,32,62^. Because I assume synNotch to be in excess, S_SJR_ can be calculated from the amount of cognate ligand a focal cell is exposed to at the given timestep. Then, S_SJR_ for a focal cell σ can be calculated using the equation

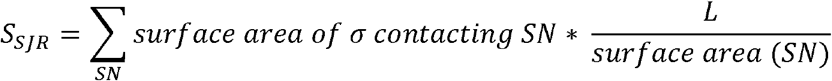

This formula is from the original GJSM study ^19^. For a focal cell σ, it receives SJR signal S_SJR_ calculated from the total number of cognate ligand it is exposed at a given timestep. For each signaling neighbor (SN) with the cognate ligand, L is the amount of cognate ligand on its surface. Dividing L by the SN’s surface area yields the ligand density. Multiplying this density by the shared surface area between the focal cell and SN gives the amount of ligand the focal cell sees from that single SN. Summing over all SNs then gives the total number of cognate ligand the focal cell is exposed to at a given timestep and yields S_SJR_.

L is a constant value for a constitutive ligand like orange ligand, but for a ligand that is inhibited by the signaling circuit (i.e. blue ligand), L is defined by R in the circuit equations. Parameters are given in Supplementary Table 1.

### Linking expression and repression to behavior

I linked expression and repression to intended behavior using a discrete transition model as in the original and accompanying study ^18,19^. Cells with protein product level exceeding the threshold (7000 for all simulations in this study) gained the feature of the protein product. For example, exceeding 7000 for E-cadherin levels allows cells to be adhesive to other E-cadherin expressing cells. Falling under this threshold results in cells losing the feature. For example, E-cadherin levels falling under 7000 reverts cells to non-adhesive.

### General simulation conditions

Cells were initialized as a 5x5x5 pixel cube in a 100x100x100 lattice. For the cell-cell signaling setup, initial configuration is specified in the results section. For the building multicellular structure experiments, cells were seeded as a spherical blob at the center of the lattice. The seeded number of cells in each blob is given per experiment. Data was collected every 100 timesteps for analysis. Simulation runtime is given per experiment.

### Cell-cell signaling assay

Assay was performed as described in the results section and in the accompanying study ^18^. Parameters are given in Supplementary Table 1.

### Structure images

Representative cross sections of the structures are shown as in the reference *in vitro* experiment^10^. Scalebar is 17.5 pixels to 100 um as determined in the original study ^19^.

### Cell percentage quantification

Cell percentage was calculated by dividing the number of cells of the focal type over total number of cells in the simulation at that timestep.

### Mitosis removal

I removed cell growth and mitosis in oscillation simulations with runtime >100000 timesteps. The longer runtime required for observing oscillation along with its termination would otherwise result in cells completely filling the lattice and slowing the simulation significantly. I validated that growth and mitosis removal did not affect oscillation (compare Fig.5C to SFig.5A, Fig.5D to SFig.5C). Cells sharply transitioned at the same timepoints and transitioned to the correct phase independent of growth and mitosis.

### Comparing oscillation quality

To roughly compare oscillation quality between circuits, I calculated the quality factor Q for each oscillation using the ring-down method ^44^. Q, the quality of the oscillation, can be defined proportionate to the number of oscillations before the oscillation decays to 50% of its maximum amplitude. I used 100% as the maximum amplitude and chose to calculate Q off the cyan phase of the oscillation as it is present in all oscillators and does not have constant drift. Q was then compared between circuits to calculate the fold improvement.

### Statistical analysis

Sample size is given in the text, figures, or captions. Plots are mean±SEM.

## SUPPLEMENTARY FIGURES

**Supplementary Figure 1.**
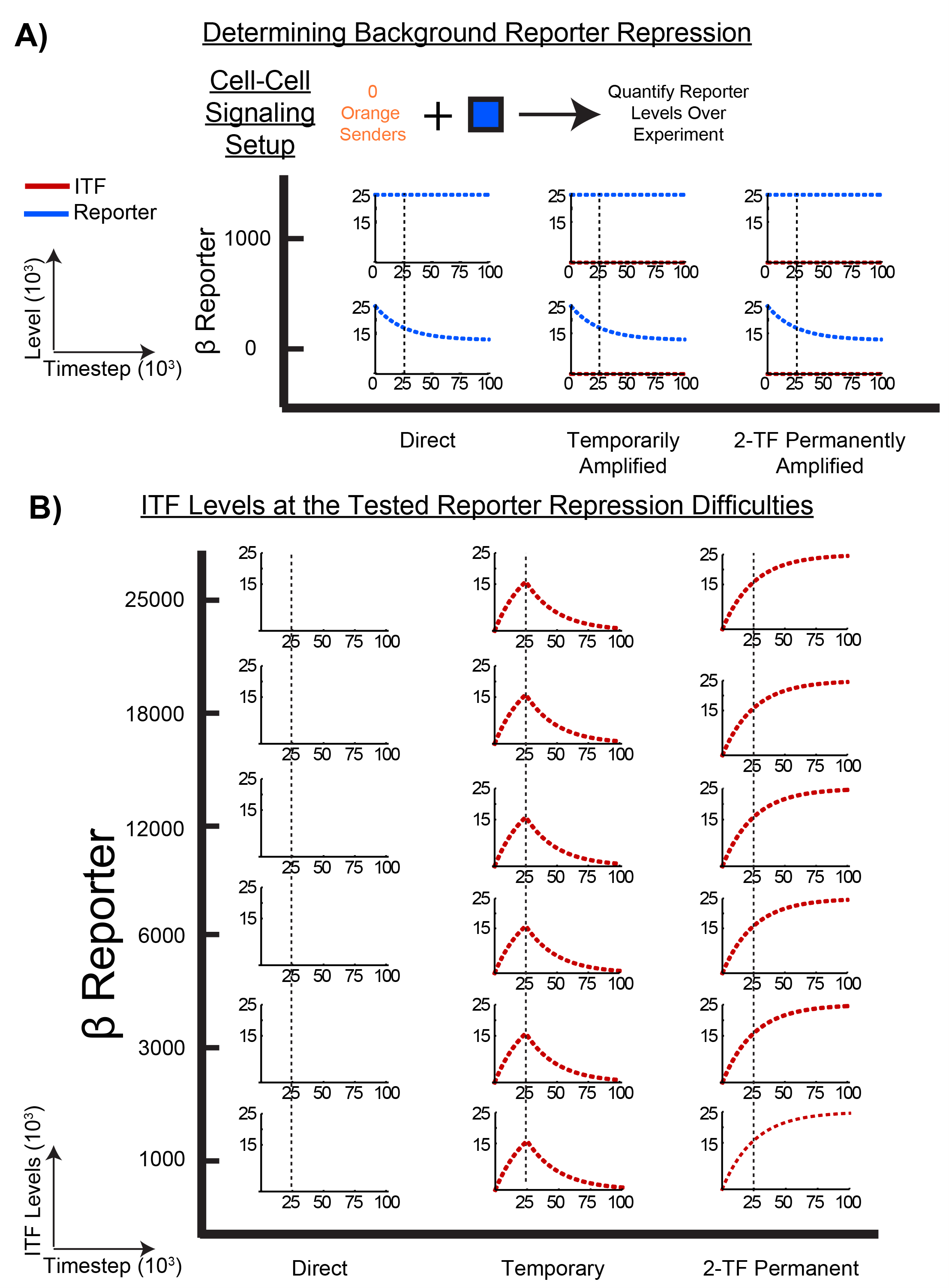
Inhibitory circuits operate as designed, related to figure 2. A) Uses the cell-cell signaling setup of Fig.2B, but blue cells are seeded without orange neighbors to determine background reporter repression. Reporter repression difficulties ≥1000 do not have leaky repression (β reporter must be at least 1000 in simulations). B) Inhibitory transcription factor (ITF) traces from the blue reporter traces of Fig.2C. One trace shown per condition. Simulations run for 100000 timesteps.

**Supplementary Figure 2.**
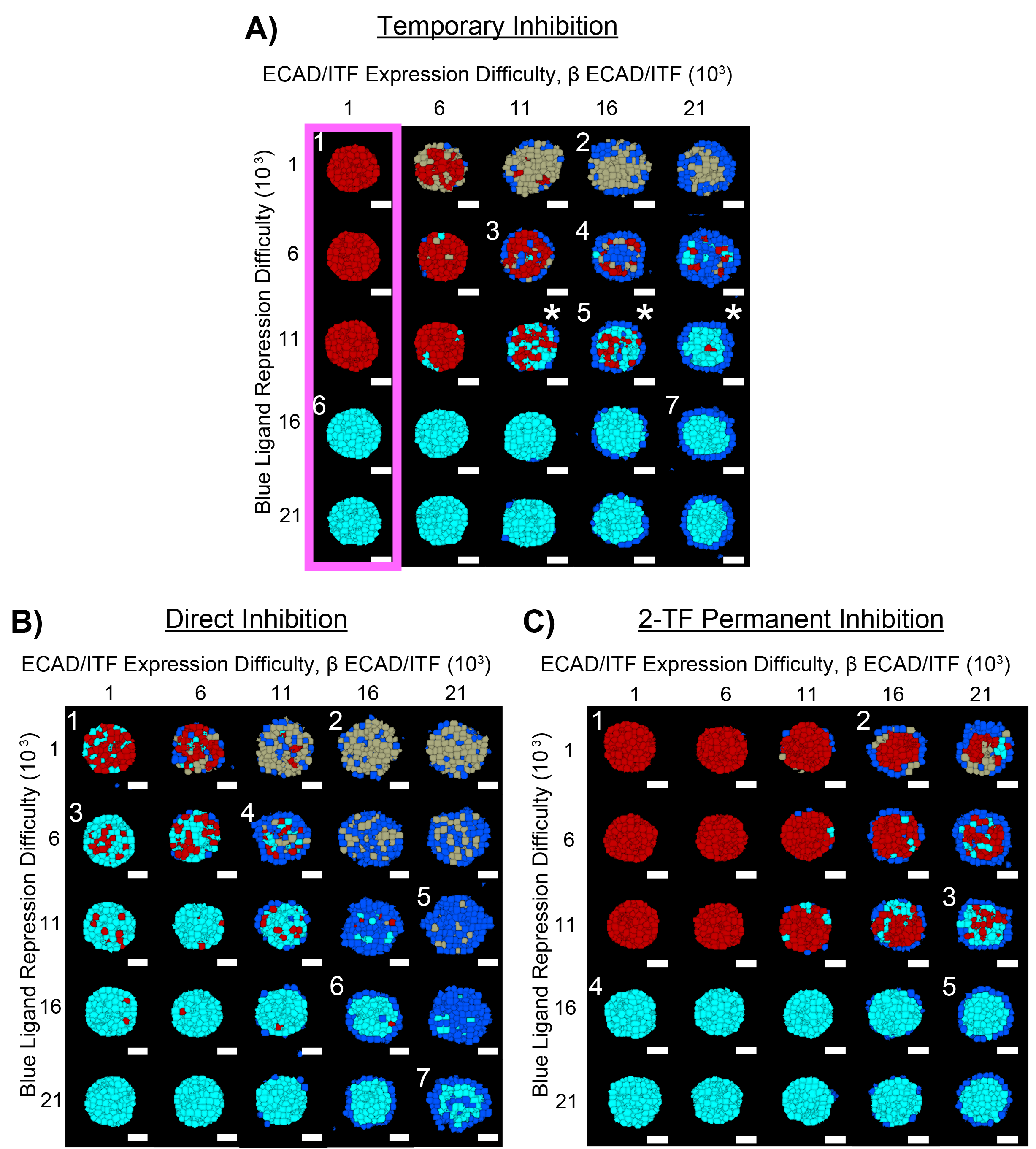
Full gallery of multicellular structures from the inhibitory circuits, related to figures 3 And 4. A) Full gallery of the structures built by the temporary inhibition circuit with parameters specified. Number labels correspond to structures shown in Fig.3E. Pink box encloses structures tested for oscillation later in the study. B) Full gallery of the structures built by the direct inhibition circuit with parameters specified. Number labels correspond to structures shown in Fig.4A. C) Full gallery of the structures built by the 2-TF permanent inhibition circuit with parameters specified. Number labels correspond to structures shown in Fig.4B. Initial mixture is 93 blue cells. One representative structure for each parameter combination is shown. N=3 for each structure. Simulations run for 50000 timesteps.

**Supplementary Figure 3.**
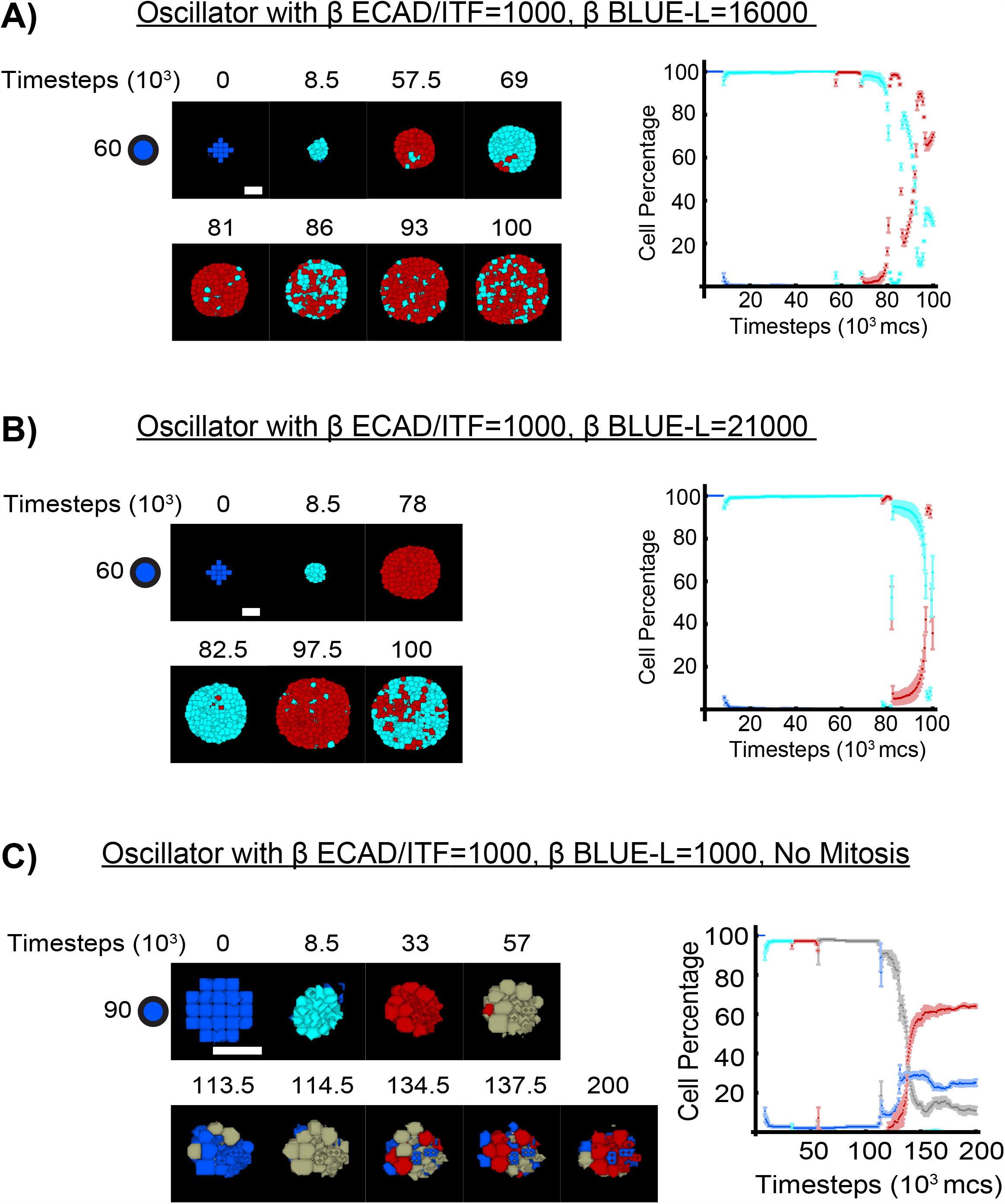
Additional oscillatory structures built by the temporary inhibition circuit, related to figure 5. A) Oscillator resulting from 57 blue cells programmed with the temporary inhibition circuit and ECAD/ITF expression difficulty=1000 (β ECAD/ITF=1000) with blue ligand repression difficulty=16000 (β BLUE-L=16000). Plot of cell percentage is given on the right. B) Same as A but with blue ligand repression difficulty=21000 (β BLUE-L=21000). C) 93 blue cells programmed with the temporary inhibition circuit and ECAD/ITF expression difficulty=1000 (β ECAD/ITF=1000) with blue ligand repression difficulty=1000 (β BLUE-L=1000) yielded poor oscillation. Mitosis was removed so the simulation could be run longer while limiting computational strain. Mitosis does not affect oscillation behavior. See Methods Section for validation. A representative developmental trajectory with images at the transition time is given per oscillation. N=3 for each oscillation. Simulations run for timesteps shown.

**Supplementary Figure 4.**
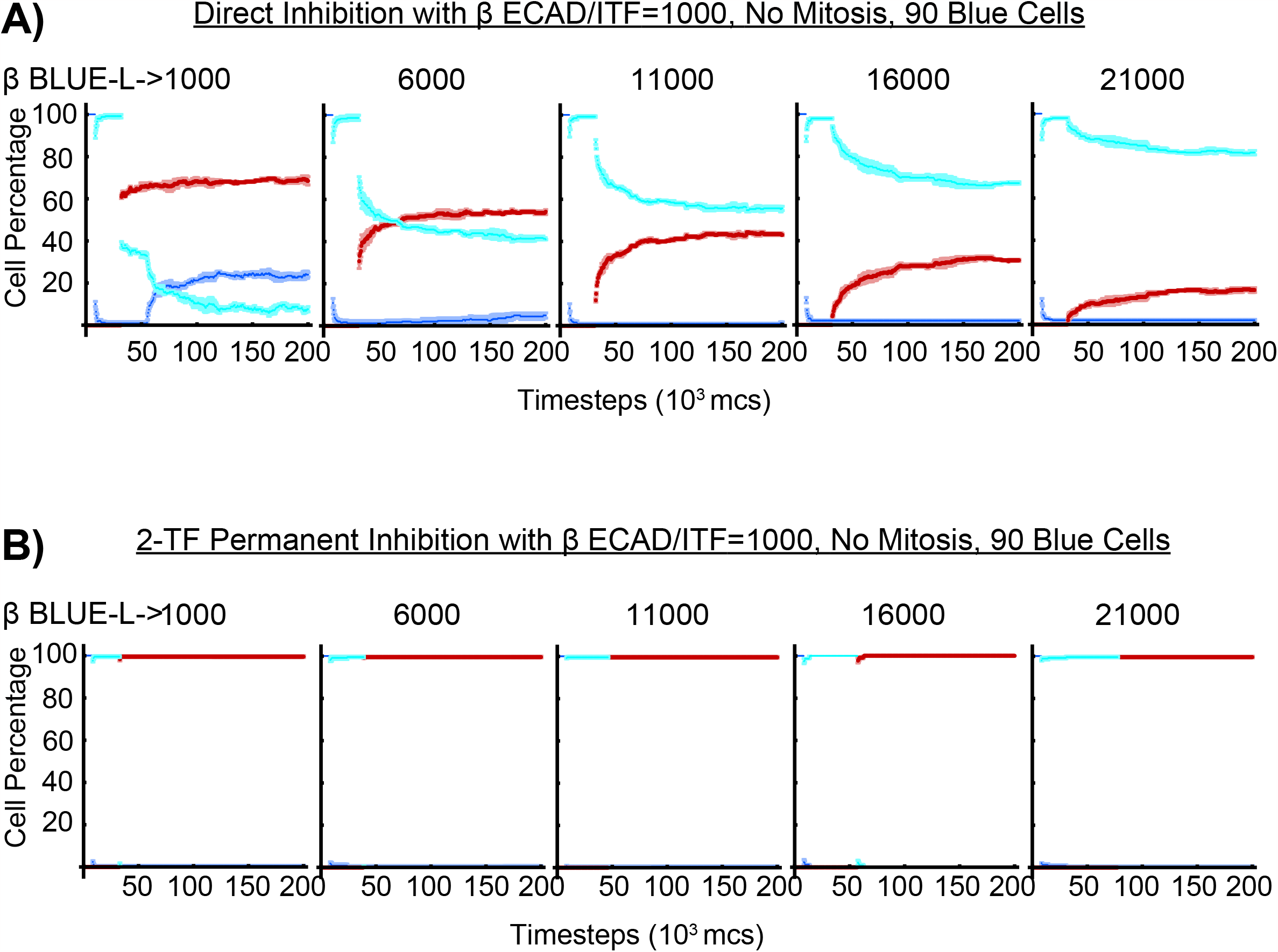
Direct inhibition circuit and 2-TF permanent inhibition circuit do not build oscillatory structures, related to figure 5. A) 93 blue cells programmed with the direct inhibition circuit were run with the same parameter combinations tested for oscillation in the temporary inhibition circuit of Fig.5 and SFig.3. There were no oscillations. B) Same as A but with the 2-TF permanent inhibition circuit. Mitosis was removed so the simulation could be run for 200000 timesteps. Plots of cell percentage is given. N=3.

**Supplementary Figure 5.**
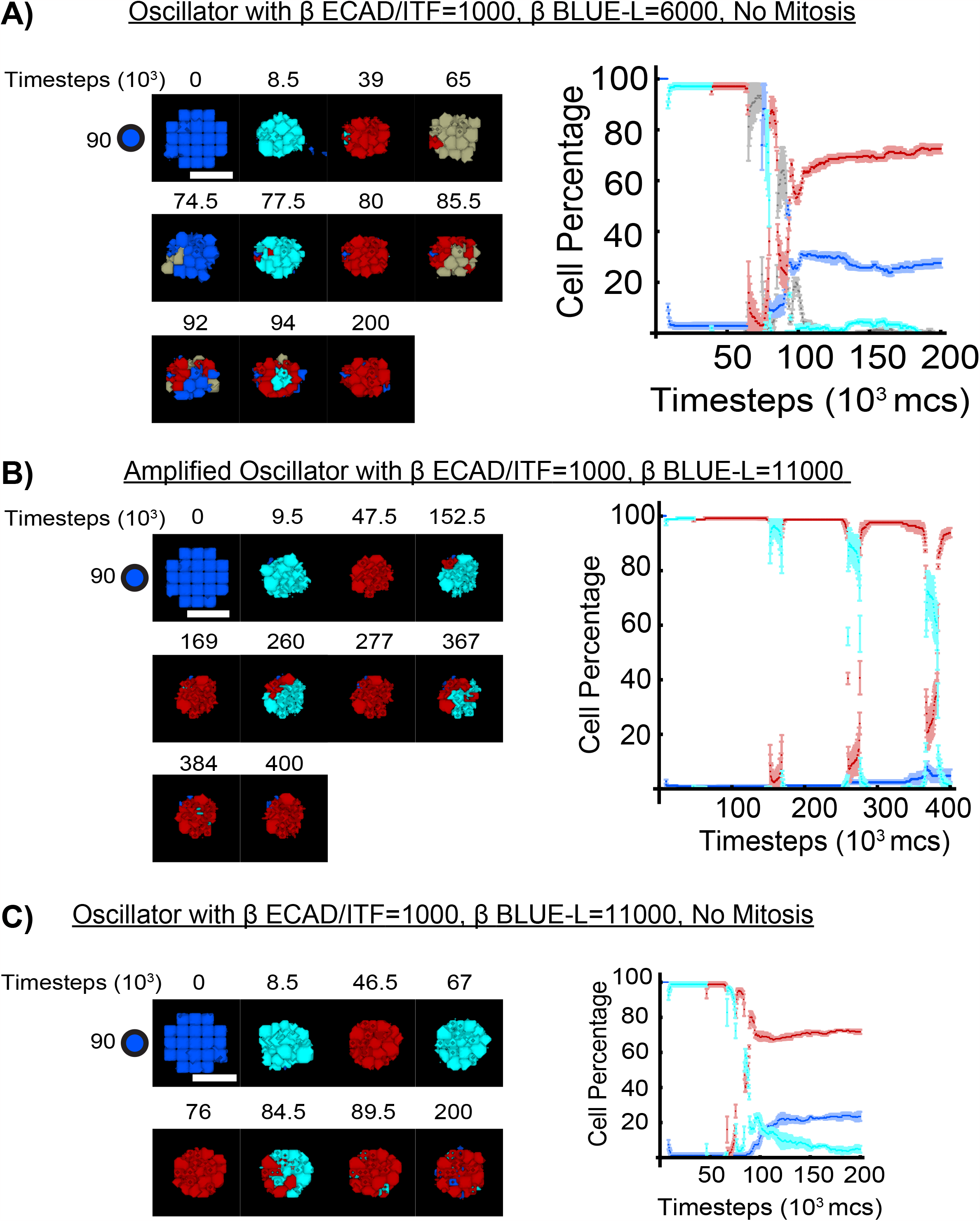
Comparing oscillatory structures built by the amplified oscillator to the oscillator (temporary inhibition circuit) with β ECAD/ITF=1000, related to figure 6. A) Oscillator resulting from 93 blue cells programmed with the temporary inhibition circuit (oscillator) and ECAD/ITF expression difficulty=1000 (β ECAD/ITF=1000) with blue ligand repression difficulty=6000 (β BLUE-L=6000). Plot of cell percentage is given on the right. Oscillation is to be compared to oscillation of Fig.6C. B) Oscillator resulting from 93 blue cells programmed with an activating amplifier controlling the temporary inhibition circuit. ECAD/ITF expression difficulty=1000 (β ECAD/ITF=1000) with blue ligand repression difficulty=11000 (β BLUE-L=11000). Plot of cell percentage is given on the right. Oscillation is to be compared to oscillation of SFig.5C (below). C) Oscillator resulting from 93 blue cells programmed with the temporary inhibition circuit (oscillator) and ECAD/ITF expression difficulty=1000 (β ECAD/ITF=1000) with blue ligand repression difficulty=11000 (β BLUE-L=11000). Plot of cell percentage is given on the right. Oscillation is to be compared to oscillation of SFig.5B. Mitosis was removed in all simulations shown here. A representative developmental trajectory with images at the transition time is given per oscillation. N=3 for each oscillation. Simulations run for timesteps shown.

**Supplementary Figure 6.**
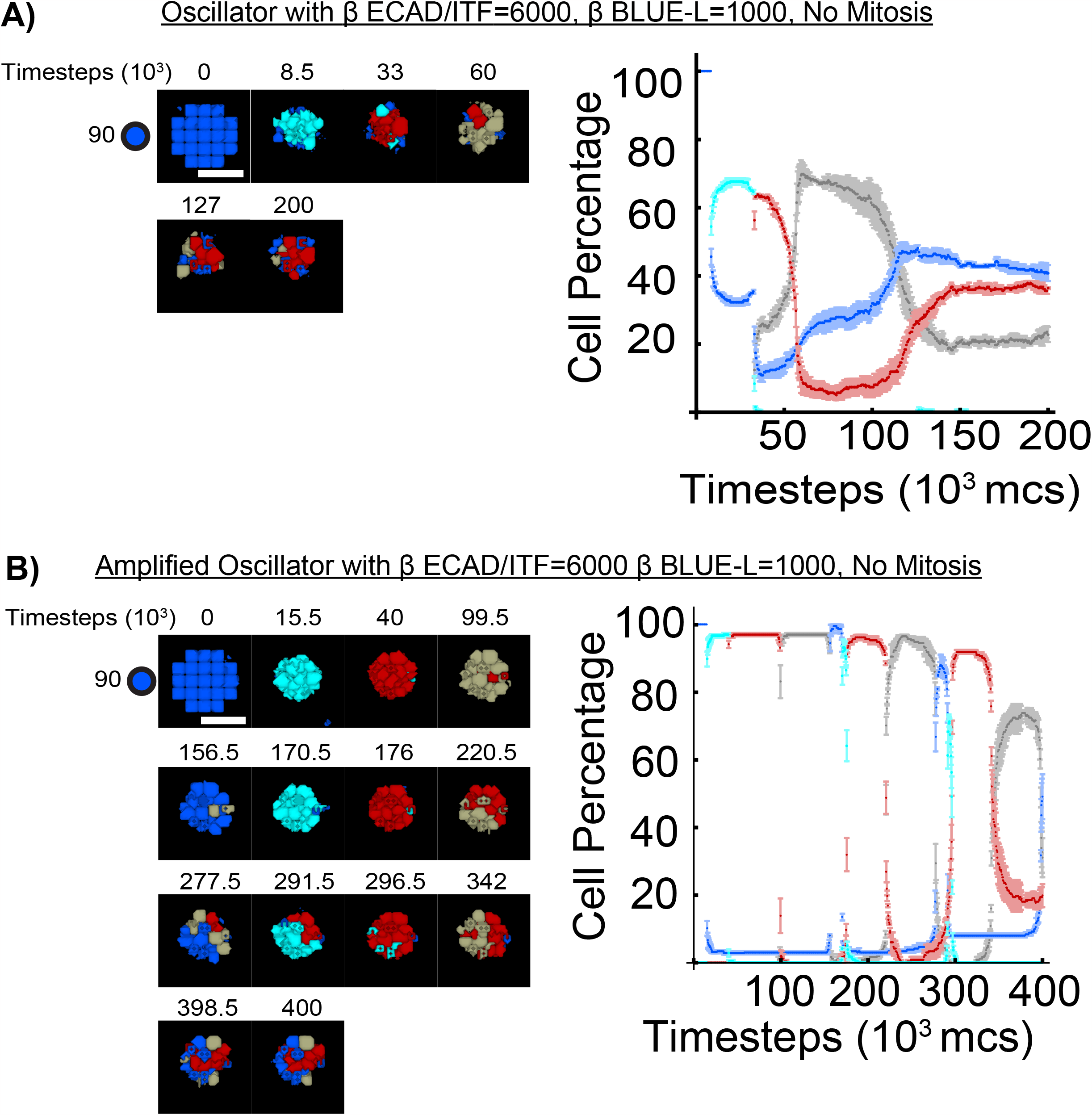
Comparing oscillatory structures built by the amplified oscillator to the oscillator (temporary inhibition circuit) with β ECAD/ITF=6000, β BLUE-L=1000, related to figure 6. A) Oscillator resulting from 93 blue cells programmed with the temporary inhibition circuit (oscillator) at given parameters. Oscillation is to be compared to oscillation of SFig.6B. B) Oscillator resulting from 93 blue cells programmed with an activating amplifier controlling the temporary inhibition circuit at given parameters. Oscillation is to be compared to oscillation of SFig.6A. Mitosis was removed in all simulations shown here. A representative developmental trajectory with images at the transition time is given per oscillation. N=3 for each oscillation. Simulations run for timesteps shown.

**Supplementary Figure 7.**
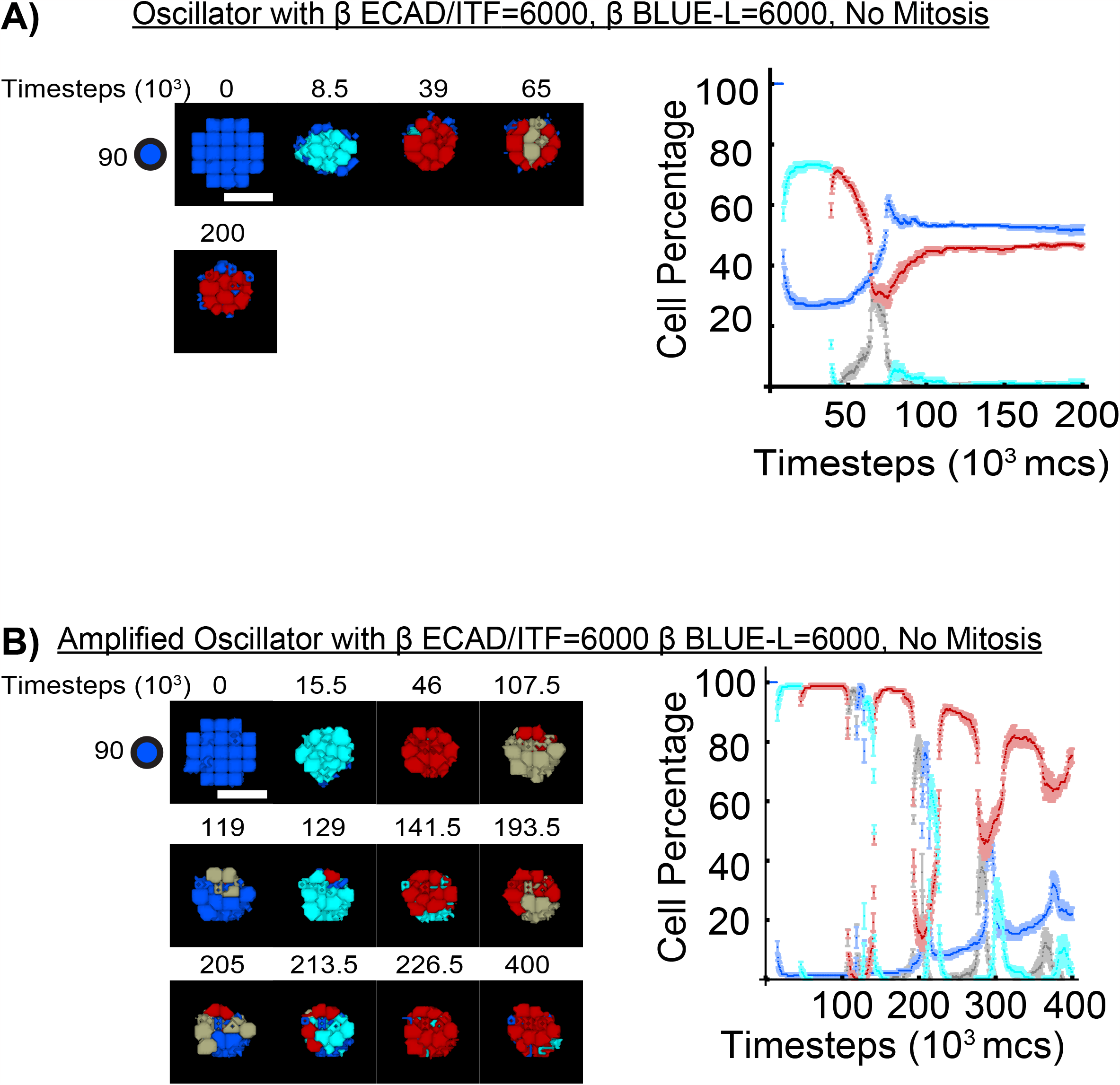
Comparing oscillatory structures built by the amplified oscillator to the oscillator (temporary inhibition circuit) with β ECAD/ITF=6000, β BLUE-L=6000, related to figure 6. A) Oscillator resulting from 93 blue cells programmed with the temporary inhibition circuit (oscillator) at given parameters. Oscillation is to be compared to oscillation of SFig.7B. B) Oscillator resulting from 93 blue cells programmed with an activating amplifier controlling the temporary inhibition circuit at given parameters. Oscillation is to be compared to oscillation of SFig.7A. Mitosis was removed in all simulations shown here. A representative developmental trajectory with images at the transition time is given per oscillation. N=3 for each oscillation. Simulations run for timesteps shown.

